# Efficient whole genome haplotyping and high-throughput single molecule phasing with barcode-linked reads

**DOI:** 10.1101/356121

**Authors:** David Redin, Tobias Frick, Hooman Aghelpasand, Jennifer Theland, Max Käller, Erik Borgström, Remi-Andre Olsen, Afshin Ahmadian

## Abstract

The future of human genomics is one that seeks to resolve the entirety of genetic variation through sequencing. The prospect of utilizing genomics for medical purposes require cost-efficient and accurate base calling, long-range haplotyping capability, and reliable calling of structural variants. Short read sequencing has lead the development towards such a future but has struggled to meet the latter two of these needs^1^. To address this limitation, we developed a technology that preserves the molecular origin of short sequencing reads, with an insignificant increase to sequencing costs. We demonstrate a novel library preparation method which enables whole genome haplotyping, long-range phasing of single DNA molecules, and *de novo* genome assembly through barcode-linked reads (BLR). Millions of random barcodes are used to reconstruct megabase-scale phase blocks and call structural variants. We also highlight the versatility of our technology by generating libraries from different organisms using only picograms to nanograms of input material.

## INTRODUCTION

Elucidating the true impact of genetic variation and its potential contribution to healthcare requires a complete characterization of the human genome. While massive efforts and technological developments have been made towards this goal, the vast majority of whole genome sequencing data produced has been limited to profiles of unphased nucleotide variations. High-throughput short read sequencing has become the backbone of genomics as a field, yet generating a haploid consensus rather than a haplotype-resolved genome has limited our ability to associate genetic variation with health and disease^1^. Deriving phenotypes from more than just profiles of SNVs (single nucleotide variations), by investigating structural variants, gene fusion events, and the cumulative effects of mutations across long distances is likely to be greatly beneficial. It is estimated that more than half of human genomic variation is constituted by structural variants^2,3^ in the form of deletions, insertions, inversions, duplications and translocations, and studies have shown such events to have a larger effect on gene expression than SNVs^4^. Identifying such variants all but equates to a need for long-range haplotype information so differences between maternal and paternal alleles can be resolved^3,5^. The importance of long-range phasing information is exemplified by the characterization of compound heterozygosity being essential for diagnosis of recessive Mendelian diseases. Furthermore, delving deeper into the genetic basis of elaborate phenotypes have shown structural variants to be drivers of cancers and complex diseases^6,7^.

Despite the promise of new-found insights for medical genomics, the adoption of whole genome haplotyping technologies have been limited by high costs due to specialized instruments and platform-dependant reagents. Assays based on dilution of genomic fragments in discrete compartments, combined with compartment-specific barcoding, have been proven effective for obtaining long-range phasing information by linking reads that share a common barcode^8–11^. Such technologies are able to utilize the high throughput and accuracy of short read sequencing platforms while maintaining the long-range information crucial for haplotyping. One technology in particular^8^, has in recent years established itself as the foremost alternative for whole genome haplotyping, but its reliance on microfluidic equipment and barcoded beads have limited its scalability and flexibility. Furthermore, the high cost of preparing libraries for sequencing has notably limited widespread adoption of this technology. Sequencing based on ‘contiguity preserving transposition on beads’^12^, offers an alternative solution for genome-wide haplotyping with a single tube reaction setup. Performing tagmentation of genomic fragments on uniquely barcoded beads, rather than in discrete compartments, opens up the potential for automated library preparation in the future. However, an evident bottleneck of this technology is the laborious generation of a transposase-linked bead library with barcodes of sufficient complexity to resolve a human genome. Furthermore, a product that enables phasing of DNA molecules has not yet been made commercially available. Alternative platforms based on long read sequencing of single molecules^13,14^ provide long-range phasing information, albeit with a lower sequencing accuracy and throughput than short read sequencing, rendering them unfit for the scale required to haplotype human-sized genomes^15–17^. Ultimately, these platforms have mostly been used to provide longer scaffolding information for genome assembly, whilst still relying on short read sequencing for coverage^18,19^.

The key to making an assay affordable is minimizing the dependency on proprietary technologies in terms of instruments and reagent kits. A number of droplet-based barcoding strategies have been developed to enable high-throughput analysis of single molecules^11^ or single cells^20–22^, all utilizing microfluidic devices for droplet generation and/or uniquely barcoded beads to distinguish the contents in each droplet. In-house manufacturing of microfluidic systems has been the solution for many academic groups to avoid commercial devices for droplet generation which are expensive and typically constricted to particular assays. Although the hardware components for microfluidic systems are easy to obtain, the manufacturing of singleuse microfluidic chips requires expertise and equipment that goes beyond what is available in most laboratories^11^. Likewise, the production of barcoded beads is not a trivial matter as highly complex libraries of unique barcodes are required to perform high-throughput analyses. The process of barcoding beads can for instance be done by clonal amplification in droplets^23^, by combinatorial extension cycling^21^ or by split-pool cycles of phosphoramidite synthesis^22^; but regardless of strategy it remains a costly and laborious pre-requisite for the intended assay reaction.

Here we describe a low-cost assay for whole genome haplotyping and single molecule phasing, based on previous work of barcoding of long DNA fragments in emulsion droplets formed by simple shaking^24^. It does not require microfluidic devices or complex libraries of barcoded beads, making it more scalable and affordable than alternative methods. Being free from these requirements also means libraries can be prepared in any laboratory setting, using readily available reagents at a cost of $19 per library and without investments in platform-specific laboratory equipment. The number of genomic fragments per droplet can be tuned according to sample input, from picograms to nanograms of DNA input, enabling haplotyping of whole human genomes, or smaller genomes with single molecule resolution. In this study, we present a haplotype-resolved human genome and investigate use of barcode-linked reads for the reference-free assembly of human and bacterial genomes.

First, a tagmentation reaction introduces a universal DNA sequence at arbitrary yet evenly distributed positions throughout the genome. Bead-linked transposases preserve the proximity of tagmented constituents from long DNA fragments, and by linking the template molecule(s) from each bead to a mutually exclusive barcode it enables the information of proximity to be conserved through DNA sequencing (**Fig. 1**). This is achieved by separating the beads into millions of discrete compartments (emulsion droplets) together with single copies of a barcoding oligonucleotide. The barcoding oligonucleotide features a semi-randomized sequence with an unrestricted complexity, ensuring the barcode present in each compartment is unique. Within each droplet, PCR amplification is used to first generate clonal populations of single stranded barcoding oligos, and then to couple the barcode to template molecules (**Supplementary Fig. 1**). As a consequence of limited dilution, droplets without either the barcoding or template molecules will be formed, but neither will yield coupled amplicons. Such products are removed from the library through a target-specific enrichment following breakage of the emulsion reaction. Libraries consisting of barcode-linked molecules are then sequenced using the standard Illumina short read platform where reads are grouped according to the barcode to reconstruct long-range haplotype information of the original fragment(s).

**Figure 1.**
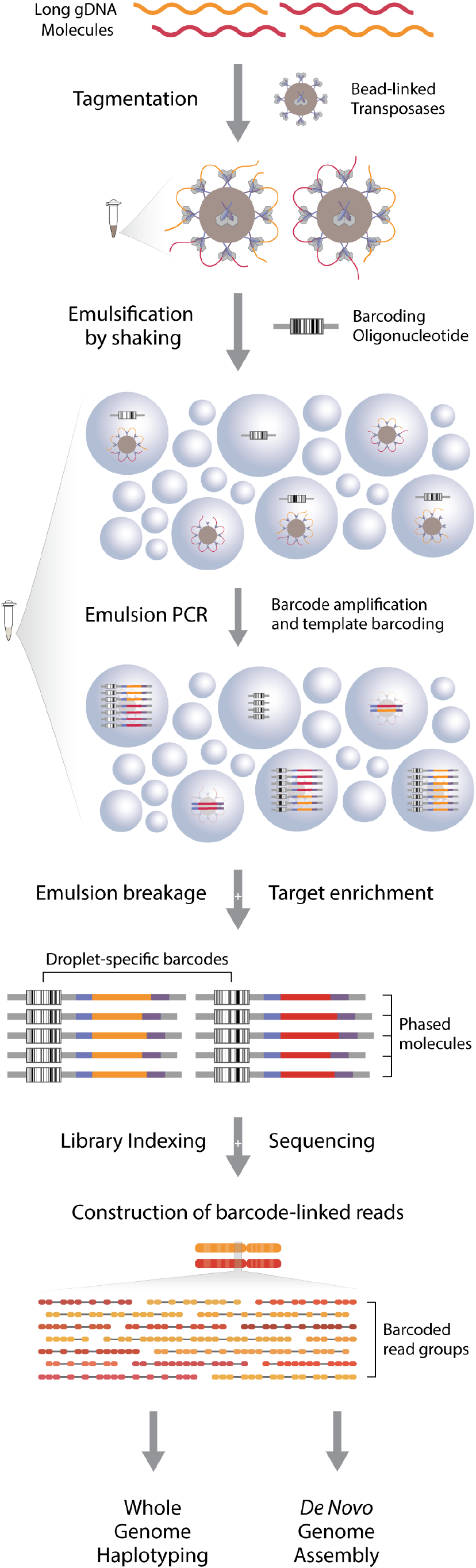
Overview of the BLR technology. High molecular weight DNA fragments are diluted and tagmented with bead-linked transposases. DNA-loaded beads are put into emulsion droplets with barcoding oligonucleotides and primers for amplification, and the constituents of each original molecule is coupled to a unique barcode sequence through emulsion PCR. Following the removal of uncoupled molecules, the library undergoes standard short read sequencing and subsequent grouping of reads according to the barcode sequence. The resulting barcode-linked reads are utilized for long range DNA phasing, genome-wide haplotyping or reference-free genome assembly.

## RESULTS

We generated barcode-linked read libraries from various sources and amounts of input DNA to validate the technology and showcase its flexibility. First, to highlight the aspect of single molecule phasing, a library was prepared with 25 pg of input DNA from *E. coli* strain BL21 and sequenced to a depth of 190X. Processing of 17,247,153 sequencing reads yielded 358,792 barcoded read groups and an N50 molecule length of 42,780 bp. A reference-free assembly of these reads initially yielded 190 contigs, which was reduced to 8 scaffolds when utilizing the barcode connections for scaffolding; with a single scaffold covering 84.2% of the genome (or 97.8% with two scaffolds). In contrast, assembling with conventional scaffolding, without taking the barcode links into account, yielded 131 scaffolds (**Fig. 2a**). Moreover, the NG50 value was 115,062 bp for the short read assembly compared to 3,812,784 bp for the barcode-linked assembly.

**Figure 2.**
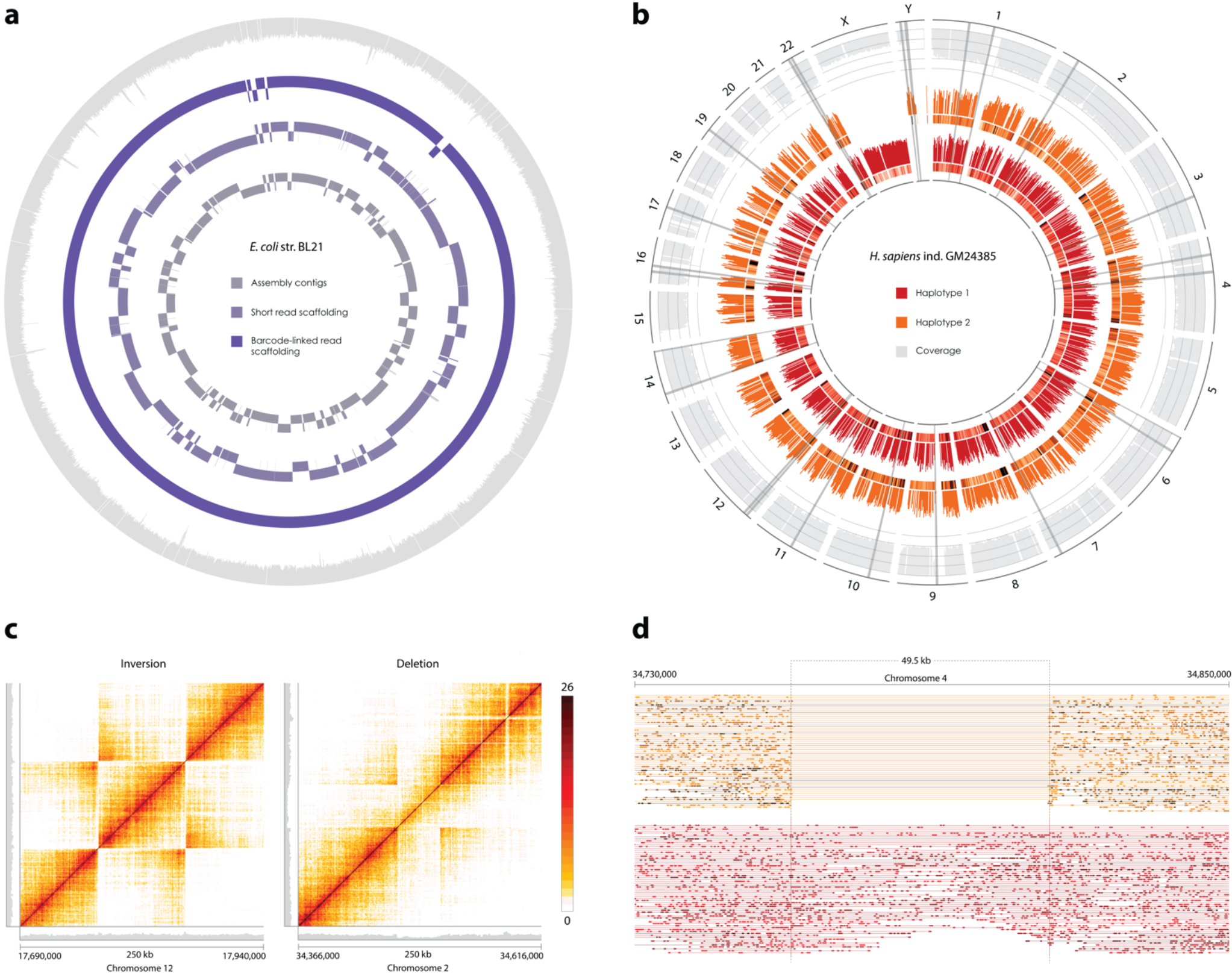
Whole genome haplotyping and de novo assembly results. (a) De novo assembly of the E. coli str. BL21 genome. From center, contigs assembled using short reads (grey), standard scaffolding performed using paired end reads (light indigo), scaffolding performed with barcode-linked reads (indigo), and the relative sequence coverage across the genome (light grey). (b) Sequence data of haplotype-resolved human genome, GM24385 (19X). From center, phased SNV density and relative read coverage for haplotype 1 (red), phased SNV density and relative read coverage for haplotype 2 (orange), total read coverage (light grey) on a scale from 0 to 25X. The localization of large structural variants are visualized by bands in grey (not drawn to scale). (c) Heatmap of barcode overlap for reads spanning called structural variants with a window size of 250 kb, an 86.0 kb inversion in chromosome 12 (left) and a 40.8 kb heterozygous deletion in chromosome 2 (right). Relative read coverage collapsed for the two haplotypes shown in grey, x and y axis are identical. (d) Barcode-linked reads of a heterozygous deletion identified in chromosome 4, with reads assigned to either haplotype, and where the reads on each line share a mutually exclusive barcode. Reads in the top haplotype (shown in orange) are linked across the deletion spanning 49.5 Kb.

Next, a library from the human ‘genome in a bottle’ (GIAB) reference individual GM24385 was generated to enable benchmarking of results against external haplotyping technologies and sequencing data. A total of 451M sequencing read pairs were processed (**Supplementary Table 1**), leaving 321M read pairs assigned to 2,204,497 barcoded read groups that were used as input for phasing analysis. With our barcode sequences translated to 10x barcodes (Methods), the Long Ranger pipeline yielded an N50 phase block length of 1,832,815 bp. The haplotype was resolved for 3,620,251 SNVs (97.9% of identified SNVs) with a mean sequencing depth of 19.1X and 0.7% of the reference genome not covered. **Figure 2b** shows that we obtain an even coverage across the whole genome and for both haplotypes, with minor discrepancies observed in some heterochromatic regions of chromosomes. The software calculated that 74.8% and 18.6% of input molecules were over 20 Kb and 100 Kb, respectively. Variant calling of constructed phase blocks yielded 29 large structural variant (LSV) calls (**Fig. 2b**) and 4,008 short deletion calls. Based on sequenced bases and mapping coordinates, we estimate the coupling efficiency of our assay to be 74.4% (**Supplementary Note**). For the purpose of benchmarking, our data was compared to a GIAB resource dataset based on the 10x Genomics Chromium Genome platform^25^, see **Supplementary Table 2** for a complete overview of output metrics from the Long Ranger analysis pipeline.

Comparing SNVs to an external ‘ground truth’ dataset^25^, based on standard Illumina sequencing to a depth of ~300X, the sensitivity of our assay was estimated to be 76.1%, with an accuracy rate of 95.4% (**Supplementary Table 3**). In context, the GIAB (10x Genomics) resource dataset with significantly higher sequencing depth, featured a sensitivity of 85.1% and accuracy of 90.7%. To exemplify the clinical relevance of phasing, our data shows all HLA family genes have their haplotypes resolved. Out of 29 large structural variant calls, 28 (96.6%) were validated through manual review of correspondence against the GIAB (10x Genomics) dataset (**Supplementary Table 4**). Called structural variants consist of inversions, deletions and duplication events across the genome, varying from 40 kb to 1.2 Mb, three of which are visualised in **Figure 2c** and **Figure 2d**.

Complementing the GM24385 library with an additional library and sequencing increased the coverage to 34.7X, enabling the evaluation of a reference-free assembly of the combined dataset. Running the human *de novo* pipeline generated an assembly with a total length of 3.2 Gb, with scaffolds covering 85.4% of the GRCh38 reference genome (heterochromatic regions not excluded, **Supplementary Fig. 2**). Scaffolding based on barcode-linked reads increased the assembly N50 from 24.4 kb to 5.60 Mb, whereof the longest contig spanned 47.9 Mb. The benefit of a higher sequencing depth was also evaluated for the Long Ranger pipeline, which with an input of 540M sequencing read pairs resulted in 98.8% of SNVs phased and phase blocks up to 11.9 megabases (N50 phase block length of 2,812,019 bp). Variant calling was in agreement with the dataset of lower sequencing depth, resulting in 30 LSVs and 4,047 short deletion calls (**Supplementary Table 2**).

## DISCUSSION

Geneticists are recognizing that short read sequencing by itself is not sufficient to resolve the connection between genetic variation and common aspects of health and disease^26^. Expanding on the capability of short read sequencing platforms to generate vast amounts of high quality data, by resolving haplotypes and calling structural variants, presents a pivotal change in genomics that enables more accurate reconstruction of genomes^27,28^. It is clear that preserving the contiguity of short sequences is currently the most affordable strategy for obtaining haplotyping information on a human genome-wide scale. We have described a novel library preparation method for barcode-linked reads that enables whole genome haplotyping for $19 per library (**Supplementary Table 5**), substantially less than the most established alternative (8). Furthermore, as the assay does not require an investment in a platform-specific instrument it can be implemented in any laboratory with a thermal cycler, in a single day (**Supplementary Fig. 3**).

Our results show that we can phase up to 99% of called SNVs in the human genome with an accuracy of ~95% and generate phase blocks with an N50 of 2.8 Mb. Analysis of data from a single lane of sequencing identified 29 large structural variants in the human genome, of which 28 could be independently verified (96.6%). Overall, the data shows that our assay is comparable to that of alternative technologies, in terms of generating long range haplotyping information, phasing SNVs (**Supplementary Table 3**) and calling LSVs (**Supplementary Table 4**) with high accuracy. To showcase the aspect of single molecule phasing and *de novo* sequencing, we also applied our method to a sample of bacterial DNA using only 25 pg of input material. The results demonstrate a drastic reduction in assembly contigs and improvement in NG50 values when barcode-linked reads are utilized compared to standard short read sequencing. To highlight the potential for *de novo* assembly of complex genomes, we were able to assemble the human genome with a modest sequencing depth of 35X using barcode-linked reads. These results showcase the power of the BLR technology for such applications.

The presented method offers laboratories all over the world the benefit of adding long-range phasing information to short read sequencing, through a simple protocol independent of specialized equipment and expensive reagents. Generating millions of discreet reaction chambers by shaking presents a convenient and scalable approach suitable for a wide range of applications. We recognize that droplets formed by shaking results in a non-uniform, albeit controllable^23^, size distribution compared to microfluidic chips. Though if microfluidic devices are readily available the proposed chemistry would be fully compatible. Regardless, the results in this study show that microfluidic devices are not necessary for producing high quality phased data. As the field seeks to draw upon the advantages of long range haplotyping information, standard practices for extracting DNA will need to shift towards more meticulous protocols aimed at maintaining the integrity of large genomic fragments for phasing. Unlike long read sequencing platforms, an advantage of our assay is that there is no inherent bias in the length of fragments that can be phased.

We present a flexible and scalable solution for whole genome haplotyping and *de novo* sequencing that can be used for DNA inputs ranging from picograms to nanograms. Consequently, it can be tailored according to the size or complexity of the genome and the resolution to which a biological study would require. An intriguing application of this technology would be reference-free assembly of complex metagenome samples, where high-throughput phasing of long single molecules is of particular interest. Furthermore, the low input requirement means haplotype-resolved genomes from single cells are feasible as a future prospect. For the purpose of expanding our understanding of population diversity and individual variance, the next frontier for large-scale genomics ought to be *de novo* and haplotype-resolved genome analyses^29^. The rise of long read sequencing and linked-read platforms show that more and more researchers are realizing the benefit of long-range phasing information. The method proposed within offers a unique opportunity for researchers to tackle the hurdles of *de novo* sequencing and genome-wide haplotyping without being limited by a lack of resources. Combined with the continued reduction in short read sequencing costs, the need for an affordable library preparation that maximizes the yield of medically relevant information is evident. In the near future, there will simply be no room for library preparation assays that cost hundreds of dollars per sample when the cost of sequencing the human genome will be reduced to a fraction of that.

## AUTHOR CONTRIBUTIONS

A.A. and D.R. conceived the technology. D.R., T.F. and A.A. designed the experiments. T.F. developed the analysis pipeline, with support from E.B., R.O and M.K.. D.R. performed all experiments, with support from H.A.. J.T. produced the bacterial *de novo* assembly. R.O. produced the human *de novo* assembly. All authors contributed in writing the manuscript.

## ACKNOWLEDGEMENTS

The authors would like to thank Christian Pou for assistance in cultivating the bacterial strain used for this study, and the National Genomics Infrastructure (NGI) and UPPMAX for providing sequencing and computational support and infrastructure.

## COMPETING INTERESTS

The authors declare no competing financial interests.

## FUNDING

This work was supported by the Erling Persson Family Foundation, and Olle Engkvist Foundation [2015/347].

## METHODS

### Sample preparation

High molecular weight gDNA was extracted from freshly cultivated cells of *E. coli* strain BL21 according to the MagAttract HMW DNA Kit (Qiagen). Following extraction the DNA was quantified by Qubit 3.0 (Life Technologies) and diluted in Elution Buffer (Qiagen). A sample of human gDNA from GIAB (Genome in a Bottle) individual GM24385 was obtained from the 10x Chromium Genome Kit (Control gDNA). On-bead tagmentation of DNA was performed using Nextera DNA Flex Library Prep (Illumina) according to the manufacturer’s reference guide for tagmentation; except that each reagent was scaled down to 15% of the specified volumes to reduce the amount of BLT (bead-linked transposases) used per reaction. The input quantity of DNA added to the on-bead tagmentation reactions varied according to the size of the genome assayed, 1 ng gDNA was used for human samples and 25 pg was used for the *E. coli* sample. Rather than amplification of tagmented DNA, the polymerase amplification was prepared without the addition of Nextera Flex indexes, and beads were subjected to an incubation at 68 degrees for 10 min instead of the specified PCR cycling protocol. The beads were subsequently washed twice with the supplied TWB reagent and then resuspended in 5 ul Elution Buffer (Qiagen) prior to emulsification.

### Reaction emulsification and retrieval

Assay reactions consist of 50 μl PCR reagents that are mixed and added on top of emulsion oil before shaking for emulsification. The PCR volumes consist of 5 ul beads with tagmented DNA (see section above), 1x Phusion Flex Master Mix (New England Biolabs), 1 M Betaine (Sigma Aldrich), 3 %vol DMSO (Thermo Scientific), 2 %wt PEG-6000 (Sigma Aldrich), 2 %vol Tween-20 (Sigma Aldrich), 400 nM Enrichment Oligo, 80 nM Coupling Oligo and 330 fM Barcoding Oligo (see **Supplementary Table 6** for oligonucleotide sequences; purchased from Integrated DNA Technologies). Emulsification is carried out by pipetting the 50 ul PCR volume on top of 100 μl HFE-7500 oil with 5 %(w/V) 008-Fluorosurfactant (Ran Biotechnologies) in a Qubit tube (Life Technologies) and shaking at 14.0 Hz for 8 min using a Tissuelyser instrument (Qiagen). Emulsion reactions were then left to stand upright for 15 min to settle, the excess oil was removed from the bottom and the remaining emulsion phase was transferred to a PCR tube with 60 μl FC-40 oil with 5 %(w/V) 008-Fluorosurfactant (Ran Biotechnologies) already added to it. 85 μl of mineral oil (Sigma Aldrich) was then added on top of the emulsion reactions, before placing the reaction tubes in a Mastercycler Pro S (Eppendorf) instrument for reaction cycling with the following protocol: 5 min at 95°C, 30 cycles of [95°C for 30 s - 55°C for 30 s - 72°C for 30 s], followed by 8 cycles of [95°C for 1 min - 40°C for 2 min - 72°C for 5 min, with 3% temperature ramp speed] and ending the protocol with 10 min at 72°C and holding at 12°C. Following emulsion PCR, the mineral oil was discarded with a pipette and 4 μl EDTA (100 mM) was added. The entire emulsion reaction and excess emulsion oil was transferred to a 0.5 ml tube (Eppendorf) and 100 μl 1*H*,1*H*,2*H*,2*H*-Perfluoro-1-octanol (Sigma Aldrich) was added before vortexing at maximum speed. After centrifugation for 1 min at 20,000 g, the aqueous phase was collected from the top and a magnetic rack was used to discard the beads.

### Sample enrichment and sequencing library preparation

Following retrieval of aqueous phases from emulsion reactions the library preparation was continued by a bead-based purification to remove short and uncoupled barcoding amplicons below 200 bp using sample purification beads included in the Nextera Flex kit. Biotinylated and barcode-coupled molecules were enriched for by washing and incubating the sample with 20 ul DynaBeads MyOne Streptavidin T1 beads (Life Technologies) in B&W buffer (1 M NaCl, 5 mM Tris-HCl, 500 μM EDTA) for 30 min under rotation at room temperature. The supernatant was then discarded and the beads washed twice with Elution Buffer, four times with NaOH (0.125 N), and finally two more times with Elution Buffer. An indexing PCR was then performed on the washed enrichment beads in 1x Phusion Flex Master Mix with 400 mM Indexing Oligo; using a protocol starting with 2 min at 95°C, 5 min at 55°C (with 10% ramp speed), 10 min at 72°C (with 3% ramp speed), and 1 min at 95°C. At this point, the PCR reaction was paused and placed on a magnetic rack (heated to 80°C), and the supernatant was transferred to a fresh PCR tube. The i5 Adapter Oligo (**Supplementary Table 6**) was then added to a final concentration of 400 nM to the reaction and the PCR indexing protocol was continued by running 4 cycles of [95°C for 30 s - 55°C for 30 s - 72°C for 1 min], followed by 2 min at 72°C. Reactions were then purified and samples were quantified by Qubit (Life Technologies). The samples were diluted to 2 nM and sequenced using the HiSeq X platform (Illumina) with 150 bp paired-end sequencing and an 8 bp single-end index.

### Data analysis overview

The analysis pipeline combines custom written python scripts, commonly used bioinformatics tools and three pipelines for sequencing reads linked by barcodes, used depending on the sample type and purpose as detailed below and outlined in **Supplementary Figure 4**. Samtools (v1.7)^30^ and pysam (v0.14)^30^ were used extensively in in-house developed scripts. All parts of the pipeline are available through GitHub (https://github.com/FrickTobias/BLR), and all sequencing data will be uploaded to the Sequence Read Archive (SRA) upon acceptance for publication.

### Read trimming and barcode deconvolution

For all samples, sequencing reads were initially trimmed with cutadapt (v1.16)^31^ using a similarity threshold of 20%, to remove universal handle sequences upstream of the barcode sequence, downstream of the barcode sequence, and downstream of genomic inserts (when applicable). Reads not following the structure and sequence of the universal handles were omitted from downstream analysis steps (**Supplementary Table 1**). The barcode sequence, matching a predefined design (**Supplementary Table 6**), were extracted from reads and divided in 8 subsets based on the first two bases, using bc_extract.py. These files were then individually clustered using CD-HIT-454 (v4.6)^32^ with k-mer cutoff of 0.9, to deconvolute the barcoded read groups.

### Bacterial *de novo* assembly

Bacterial samples were first required to have their barcodes grouped by making an initial assembly with IDBA-UD^33^ followed by mapping original reads to it with BWA^34^. Mappings were put into the barcode-aware Athena^35^ to yield an assembly. This assembly was preprocessed with ARCS (v1.0.1)^36^ to enable scaffolding of linked-read data with LINKS (v1.8.5)^37^, using a head tail region of 5 kb for the final assembly. The conventional short-read assembly was instead generated by scaffolding the initial non-linked assembly using SPACE (v2.2)^38^. Bacterial assembly metrics were calculated using metaQUAST (v5.0.0)^39^.

### Human genome haplotyping and *de novo* assembly

Trimmed reads were first mapped using Bowtie2 (v2.3.4.1)^40^ with GRCh38 as reference, thereafter read duplicates with the same barcode were called and removed using picard tools (v2.5.0) (http://broadinstitute.github.io/picard/). To identify and collapse barcode sequences originating from the same droplet, barcode-linked reads sharing at least two proximal (<100 Kb) read pairs were merged using cluster_rmdup.py. To exclude potentially erroneous phasing information from abnormally large droplets, the barcode sequence was stripped from all reads originating from droplets with >260 molecules using filter_clusters.py. For this purpose, molecules were defined as barcode-linked reads whereby each read mapped no further than 30 kb from its closest neighbor. Reads along with corresponding barcodes were then converted in accordance to the input of the 10x Genomics analysis pipelines (www.10xgenomics.com) with wfa2tenx.py. To enable use of these pipelines, which feature a limit of 4.8M barcodes, the complexity of our barcode population was reduced by stripping the barcode information from all barcoded read groups with less than 4 read pairs. For whole genome haplotyping analysis, the Long Ranger analysis pipeline (v2.2.2) was run with GATK (v3.8) for SNV calling and a 10x GRCh38 blacklist to reduce false positive variant calls. For the *de novo* assembly, the Supernova^41^ pipeline was utilized, with the option of no-preflight to allow for shorter insert sizes than the typical Chromium data.

### Data validation and visualization

For *de novo* assemblies, the final scaffolds zero coverage and mismatch percentages were calculated based on metrics supplied from metaQUAST (39) for the bacterial assembly and QUAST (4.6.4)^42^ for the human genome assembly respectively. Human genome scaffolds were aligned using Minimap2 (v2.4)^43^. SNV calls were evaluated by comparing output variant files from the Long Ranger analysis to variant calls from a reference dataset (300X Illumina sequencing)^25^ using VCFtools^44^ and large structural variation calls (>30kb) were compared manually (**Supplementary Table 4, Supplementary Fig. 5, Supplementary Fig. 6**) to variants identified in the resource dataset GIAB (10x Genomics)^25^. The GIAB (10x Genomics) resource dataset was downloaded and run through Long Ranger with the same version and parameters as previously described. In addition to previously mentioned softwares, MultiQC (v1.6.dev0)^45^, bedtools (v2.27.1)^46^, Circos (v0.69.6)^47^ and dotPlotly (https://github.com/tpoorten/dotPlotly) were used for calculations and data visualization.

